# Environmental modulation of global epistasis is governed by effective genetic interactions

**DOI:** 10.1101/2022.11.02.514859

**Authors:** Juan Diaz-Colunga, Alvaro Sanchez, C. Brandon Ogbunugafor

## Abstract

Interactions between mutations (*epistasis*) can add substantial complexity to genotype-phenotype maps, hampering our ability to predict evolution. Yet, recent studies have shown that the fitness effect of a mutation can often be predicted from the fitness of its genetic background using simple, linear relationships. This phenomenon, termed *global epistasis*, has been leveraged to reconstruct fitness landscapes and infer adaptive trajectories in a wide variety of contexts. However, little attention has been paid to how patterns of global epistasis may be affected by environmental variation — both from external sources or induced by the population itself through eco-evolutionary feedbacks — despite this variation frequently being a major driver of evolution. By analyzing a four-mutation fitness landscape, here we show that patterns of global epistasis can be strongly modulated by the concentration of a drug in the environment. Using previous theoretical results, we demonstrate that this modulation can be explained by how specific gene-by-gene interactions are modified by drug dose. Importantly, our results highlight the need to incorporate potential environmental variation into the global epistasis framework in order to predict adaptation in dynamic environments.

---

The topography of genotype-phenotype maps has critical consequences for the predictability of evolutionary trajectories. This topography emerges from complex interactions between genetic elements, from single nucleotides to protein residues and metabolic pathways (*1*–*5*). Despite this complexity, recent work has shown that epistasis — the nonlinear interaction between parcels of genetic information — often has a “global” component (*6*–*14*), emerging in the form of simple linear correlations between the fitness effect of a mutation and the fitness of the genetic background where it arises (Fig. 1A-C). *Global epistasis* has become a central concept in modern conversations surrounding the fundamentals of evolutionary processes. Negatively sloped correlations have been more commonly reported, both in the form of *diminishing returns* and *increasing costs* epistasis (where the fitness effect of a mutation becomes less beneficial or more deleterious, respectively, in fitter genetic backgrounds) (*6*–*8*,*11*–*13*). Positive slopes (*increasing returns* or *decreasing costs* epistasis) have also been found in low-dimensional landscapes (*7*,*15*,*16*), and theory suggests they may be more common near fitness peaks (*5*,*17*,*18*).

**Fig. 1.**
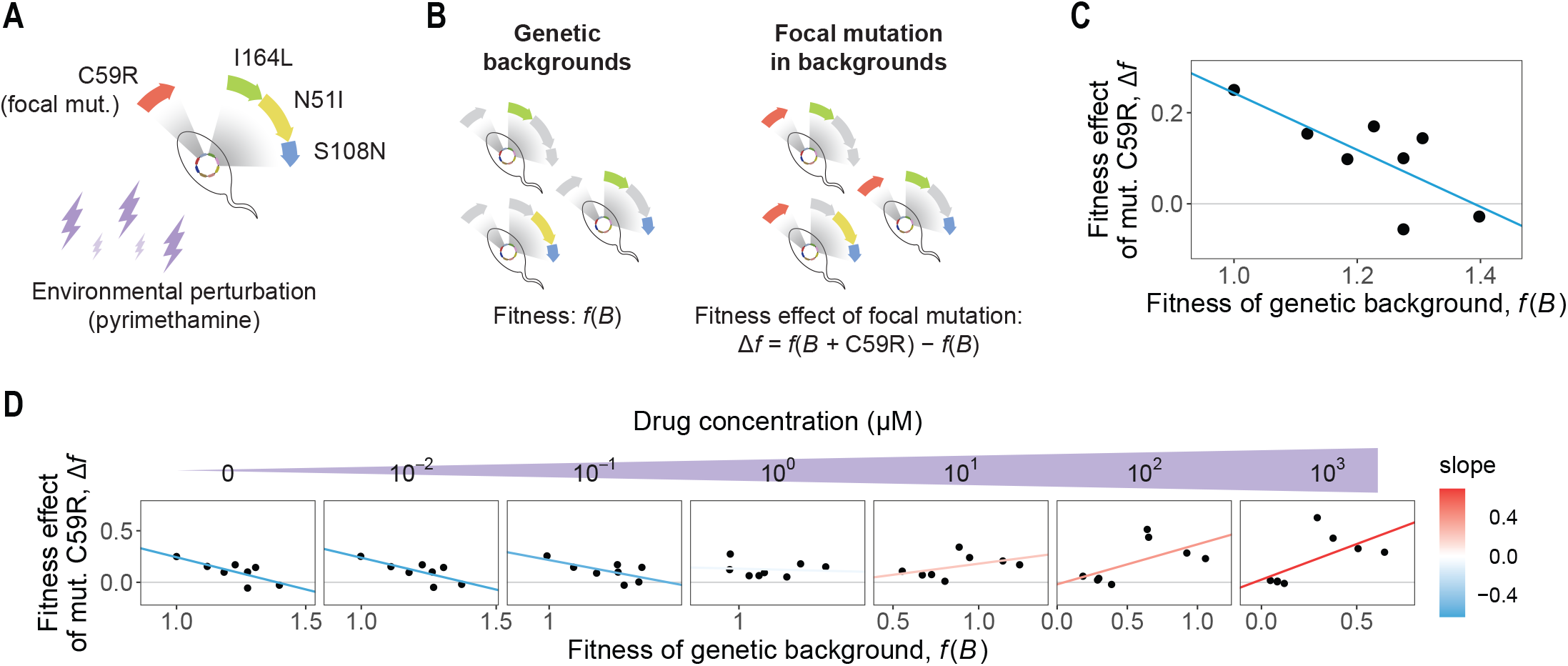
Environmental perturbations can modulate the shape of global epistasis. (**A**) We reanalyzed a dataset consisting of 16 alleles of the *P. falciparum* parasite, each carrying a different combination of four mutations: C59R, I164L, N51I and S108N. (**B**) We examined the fitness effect of the focal mutation C59R in different genetic backgrounds consisting of every possible combination of the other three mutations. In this illustration, colored/gray genes represent the presence/absence of the mutation. (**C**) In the absence of pyrimethamine, a negative correlation is observed between the fitness effect of the focal mutation and the fitness of its genetic backgrounds. (**D**) As the concentration of pyrimethamine increases in the environment, the slope of the global epistasis pattern for mutation C59R becomes more positive.

The emergence of global epistasis evidences the existence of regularities in genotype-phenotype maps (*6*–*10*), which have been leveraged in recent methodologies to infer complete adaptive landscapes under defined, steady environmental conditions (*19*–*21*). Yet, the magnitude of mutational fitness effects (*22*–*25*) and epistatic interactions (*26*–*30*), and thus the topography of fitness landscapes (*31*–*33*), can generally depend on environmental variables. Populations are often subject to natural or anthropogenic environmental fluctuations which can dictate their evolutionary fate (*25*,*28*,*34*,*35*) — this, for instance, has been widely described in the evolution of antibiotic resistance (*28*,*36*–*38*). Organisms themselves can also modify their habitat as they grow, altering downstream evolution (*33*,*39*–*41*). Learning how global epistasis patterns may be shaped by environmental factors is therefore critical to our ability to predict adaptation. This is yet an open question largely because understanding how global epistasis emerges from fine-grained genetic interactions (which might be subject to environmental regulation) is still in its early days.

Here we ask whether the environment may modulate the patterns of global epistasis observed for a particular mutation and, if so, whether we can trace back the origins of this modulation to specific gene-by-environment interactions. To address these challenges, we analyzed a previously published dataset (*30*,*32*,*42*) consisting of 16 alleles of the *P. falciparum* parasite, each carrying a different combination of four mutations: C59R, I164L, N51I and S108N (Fig. 1A). Each of the 16 genotypes was cultured in a concentration gradient of pyrimethamine or cycloguanil (antifolate drugs used to treat malaria), and fitness was quantified as the growth rate relative to that of the slowest growing allele. For convenience, in this paper we focus on C59R as the *focal* mutation, and we refer to I164L, N51I and S108N as the *non-focal* mutations (or *background* mutations). We also focus on pyrimethamine as our model environmental perturbation. Equivalent analyses for cycloguanil and for every mutation other than C59R as the focal can be found in the Supplementary Material.

Our results show that the environment can modulate the patterns of global epistasis observed in this system, which can range from *diminishing returns* to *accelerating returns* for a same mutation at different drug doses. Making use of previous theoretical results, we demonstrate that this modulation can be explained by specific gene-by-gene interactions and how they are affected by the environment. In particular, we mathematically define a set of “effective” genetic interactions, showing that they can serve to track the shape of global epistasis across drug concentrations. Finally, we turn to a simple model of epistasis to provide intuition regarding how these effective interactions relate to pairwise epistasis between mutations.

## The environment modulates the shape of global epistasis patterns

We analyzed every concentration of pyrimethamine ranging from 10^−2^ μM to 10^3^ μM, as well as the no drug control, in the dataset described above. We refer to the set of genotypes not carrying mutation C59R as the *genetic backgrounds* of this focal mutation (Fig. 1B). We denote *f* (*B*) the fitness of one of such backgrounds *B*. Calling *B*+C59R the allele resulting from adding mutation C59R to the genetic background *B*, the fitness effect of C59R is quantified simply as: Δ*f* = *f* (*B*+C59R) − *f* (*B*), that is, the difference in fitness between alleles *B*+C59R and *B* (Fig. 1B).

In the absence of pyrimethamine, we observed clear signatures of global epistasis for the focal mutation. The fitness effect of C59R negatively correlates with the fitness of its genetic background, following a pattern of *diminishing returns* commonly reported in genetic studies: the mutation becomes less beneficial in backgrounds of higher fitness (Fig. 1C). As the drug concentration increases, however, this pattern inverts and the slope becomes positive (Fig. 1D). At the highest doses, global epistasis follows a pattern of *increasing returns*, where mutation C59R becomes more beneficial in fitter genetic backgrounds. For other focal mutations, the global epistasis slope is modulated differently by pyrimethamine, for instance going from flat to negative for S108N at intermediate concentrations (fig. S1). Using cycloguanil as the environmental perturbation also changes the way in which the slopes vary with drug concentration: the focal mutation C59R, for instance, exhibits a slope that goes from negative to flat (fig. S2). This observation highlights the role of the environment in dictating the shape of global epistasis in this particular system.

### Environmental modulation of global epistasis is explained by changes in background fitness effects and interactions

Can we attribute these environmental effects on global epistasis to specific genetic interactions? Recent work has demonstrated that global epistasis patterns can emerge as a result of just a few (*14*), or multiple widespread (*43*), gene-by-gene idiosyncratic interactions. In the latter case, a recent study has found an explicit quantitative relationship connecting the shape of global epistasis to microscopic interactions between mutations (*43*). Theory has suggested that this relationship may hold even when such interactions are sparse as opposed to widespread (*44*). In particular, these works show that the global epistasis slope (*b*_C_) of a focal mutation (here C59R, denoted as C) can be approximated by a sum of contributions from every other mutation (*i* ≠ C, that is, *i* = I164L, N51I or S108N) as (*43*,*44*):

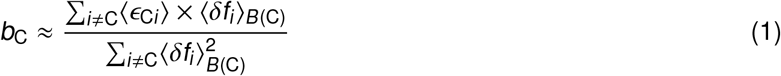

The term ⟨δ*f_i_*⟩_*B*(C)_ represents the average fitness effect of mutation *i* across all backgrounds where mutation C59R may be introduced, that is, all genotypes not carrying C59R. This set of backgrounds is denoted as *B*(C). The term ⟨*ϵ*_C*i*_⟩ represents the average pairwise epistasis between mutations *i* and C59R, defined as the deviation between the fitness of the double mutant with respect to the expectation that mutations *i* and C59R act additively. This average is taken across every possible background in which mutations *i* and C59R may be introduced, i.e., every allele not carrying any of the two. Fig. 2A-B provides a simple intuitive interpretation for all the magnitudes defined above. Equation 1 can be expressed simply as

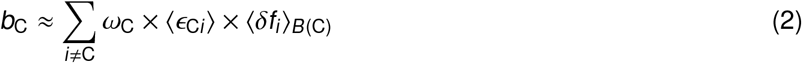

Where we have defined the “weight” *ω*_C_ as just

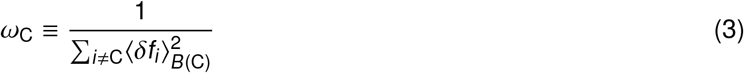

Note that *ω*_C_ is always positive, and therefore equation 2 indicates that the sign of the slope results from (a) the magnitude and sign of the fitness effects of every background mutation (captured by the terms ⟨δ*f_i_*⟩), and (b) the interactions between said non-focal mutations and the focal C59R (captured by the terms ⟨*ϵ*_C*i*_⟩). The factor *ω*_C_ × ⟨*ϵ*_C*i*_⟩ × ⟨δ*f_i_*⟩ may be interpreted as a quantification of an “effective interaction” between mutations C59R and *i*, which depends not only on the epistatic coefficients *ϵ*_C*i*_ but also on the fitness effects δ*f_i_*. Intuitively, this indicates that even if epistasis between two mutations is strong (large ⟨*ϵ*_C*i*_⟩), this will not affect the slope unless the non-focal mutation has a high fitness effect (large ⟨δ*f_i_*⟩). The sum of these effective interaction terms determines the global epistasis slope of the focal mutation, and thus this sum may be seen as a representation of a coarse-grained “mutation-by-background” (or “gene-by-genome”) interaction. In light of equation 2, we may expect environmental factors to modulate the shape of global epistasis through two different mechanisms: either by altering the fitness effects of the background mutations, or by modulating the interactions between them and C59R (Fig. 2C).

**Fig. 2.**
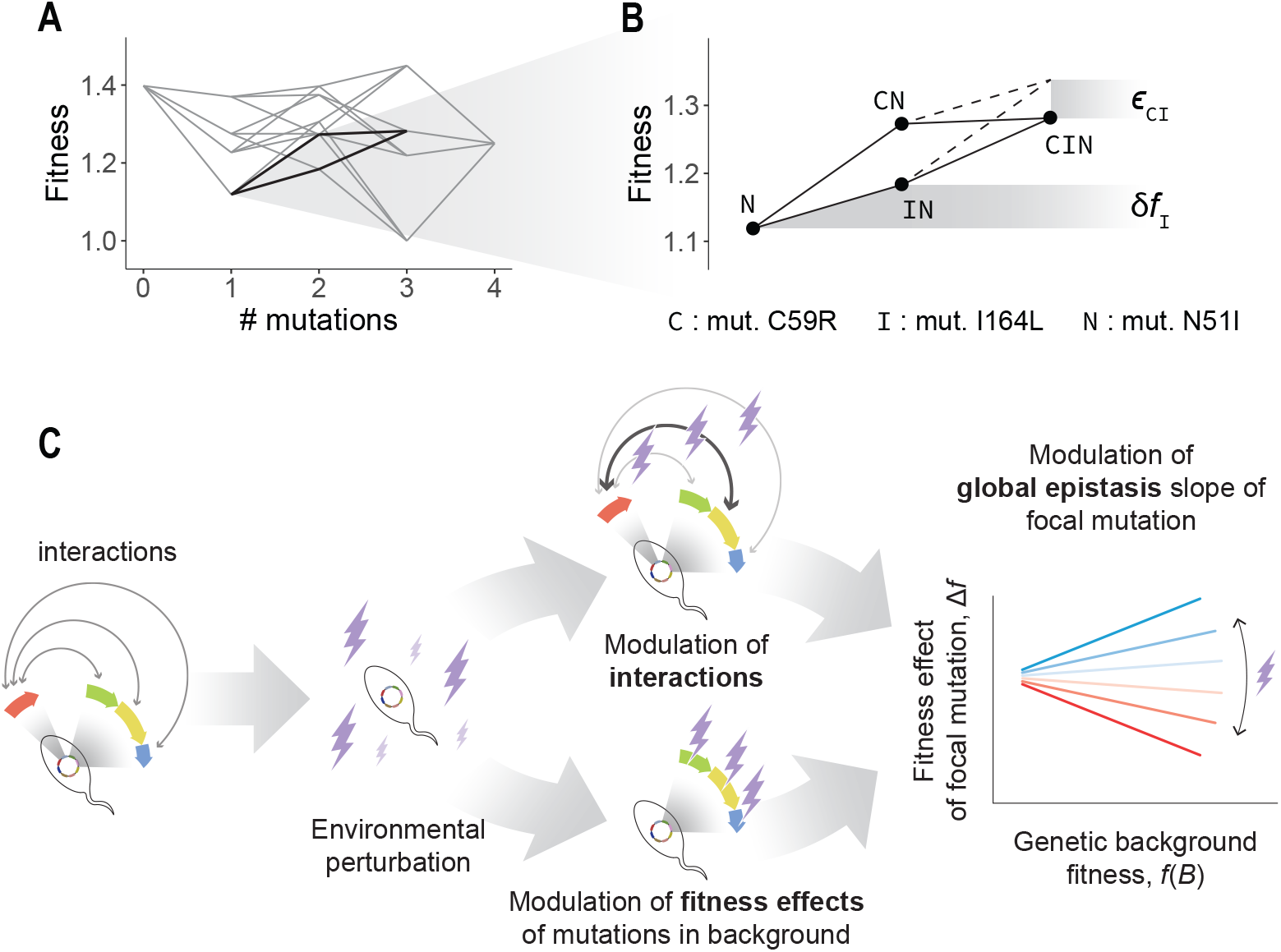
Potential mechanisms for the environmental modulation of global epistasis. (**A**) Fitness graph of the system analyzed here. Each node corresponds to an alleles, edges connect genotypes at a distance of one mutation. (**B**) As an example, we show the detail of four specific alleles: the genetic background carrying mutation N51I only, and the three genotypes resulting from the addition of mutation I164L, C59R, or both to the background. The fitness effect of I164L is quantified as the difference in fitness between the allele carrying both I164L and N51I, and the background carrying N51I only. The epistatic interaction between C59R and I164L is quantified as the deviation of the triple mutant with respect to the additive expectation (dashed lines), and is negative in this case (*ϵ*_CI_ < 0). (**C**) Environmental perturbations may modulate global epistasis by affecting either the fitness effects of the background mutations or their interactions with the focal mutation (equations 1 and 2).

To test these two potential mechanisms, we quantified the terms δ*f_i_* and *ϵ*_C*i*_ for every non-focal mutation *i* in the dataset and for every drug concentration between 0 and 10^3^ μM. We did this in every possible background of C59R (as an example, see the case where the background is the allele carrying just mutation N51I in Fig. 2B). We observed that all three background mutations exhibit different behaviors. First, the fitness effects (*δ*) of I164L remain relatively low across all drug concentrations, and so does the magnitude of the pairwise interaction (**ϵ**) between I164L and C59R (Fig. 3A-B, green). This indicates that I164L should have only a modest contribution to the global epistasis slope of C59R. Mutation N51I exhibits a negative average fitness effect in the absence of pyrimethamine, which gradually becomes closer to zero as the drug concentration increases (Fig. 3A, yellow). Its interaction with the focal mutation C59R is positive for low drug concentrations and goes to zero for higher doses (Fig. 3B, yellow). Therefore, we can expect N51I to contribute to the global epistasis slope of C59R negatively and only for low pyrimethamine doses. Finally, mutation S108N displays the opposite behavior: its fitness effect is small for low doses but becomes large and positive at higher concentrations, and so does its interaction with C59R (Fig. 3A-B, blue). Its contribution to the global epistasis slope of C59R should thus be insignificant at low drug doses, and become positive at high doses.

**Fig. 3.**
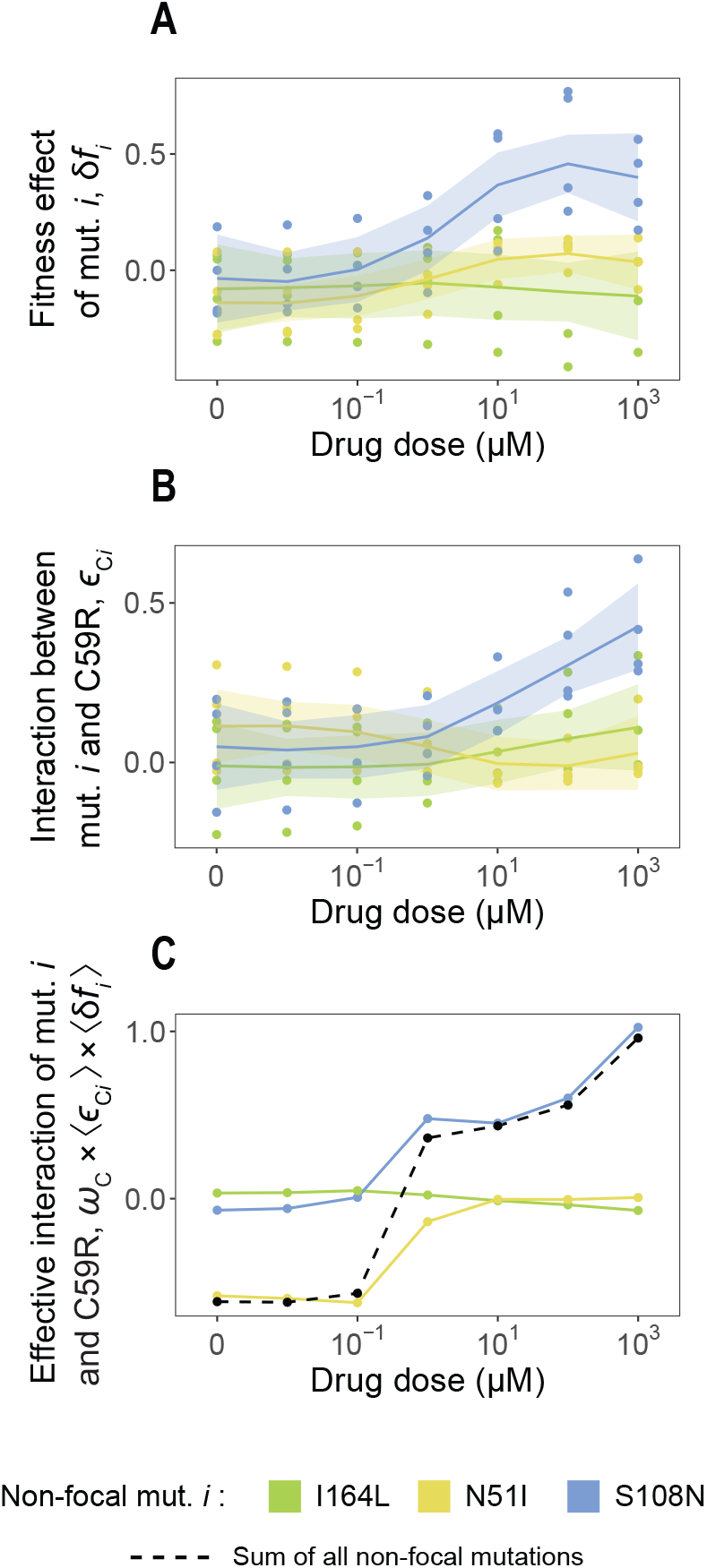
Effective interactions explain the effect of the environment on global epistasis for mutation C59R. (**A**) We quantified the fitness effect of mutations I164L (green), N51I (yellow) and S108N (blue) on each background not carrying the focal mutation C59R and for every pyrimethamine dose. Each dot corresponds to a different background. Lines and shaded regions are LOESS regressions and 95% confidence intervals of these. (**B**) In every background not carrying mutation C59R, we also quantified the magnitude of the interaction between C59R and each of the three background mutations, i.e., the deviation between the fitness of the double mutant with respect to the additive expectation (see Fig. 2B). (**C**) Effective interactions between C59R and the three background mutations are quantified as the product *ω*_C_ × ⟨*ϵ*_C*i*_⟩ × ⟨*δf_i_*⟩ (see equation 2). The sum of these products (dashed line) is dominated by the effective interaction between C59R and N51I at low pyrimethamine doses (yellow line), and by the interaction between C59R and S108N at high doses (blue line).

We tested this reasoning by quantifying the effective interactions between C59R and each of the three background mutations, i.e., the products *ω*_C_ × ⟨*ϵ*_C*i*_⟩ × ⟨*δf_i_*⟩, for every drug concentration (note that *ω*_C_ also varies with pyrimethamine dose, since it depends on the fitness effects of the non-focal mutations and these can be modulated by drug concentration, see fig. S3). As expected, the effective interactions with C59R follow the patterns described above: the term corresponding to I164L remains close to zero across the entire range of pyrimethamine doses, the term for N51I goes from negative to zero, and the term for S108N goes from zero to positive. The sum of these effective interactions (∑_i_*ω*_C_ × ⟨*ϵ*_C*i*_⟩ × ⟨*δf_i_*⟩, that is, the estimated slope *b*_C_ of the focal mutation C59R) therefore goes from negative to positive as pyrimethamine dose increases (Fig. 3C-D). When it is negative, it is dominated by the contribution of N51I, and when it becomes positive, the contribution of S108N dominates (Fig. 3C). Analyses for the patterns exhibited by other mutations are included in fig. S4 (pyrimethamine) and fig. S5 (cycloguanil).

These results highlight the utility of defining effective interactions (equation 2) in order to uncover which microscopic genetic interactions, and through which mechanisms, play a more prominent role in dictating the shape of global epistasis across environments. Our analysis shows that equations 1 and 2 can explain the global epistasis slope of C59R even when it is dominated by a single interaction (C59R-N51I at low doses and C59R-S108N at high doses), that is, in the limit of sparse interactions. This provides empirical support for recent theoretical results (*44*). Naturally, focal mutations other than C59R exhibit patterns of global epistasis that can be environmentally modulated in different ways, both by pyrimethamine or cycloguanil. For some of the other mutations, this modulation is also dominated by a single interaction (see e.g. mutation N51I in fig. S4, where the slope is dictated by its interaction with S108N). But, in other cases, global epistasis can result from the superposition of more widespread interactions (see e.g. the same mutation N51I in cycloguanil, fig S5). This observation highlights the value of effective interactions, as defined in equations 1 and 2, to understand the environmental modulation of global epistasis in a variety of contexts (figs. S1 to S5).

### A simple model of epistasis recapitulates empirical observations

To gain intuition about the mechanisms for the environmental modulation of global epistasis, we found it useful to study a simple minimal model. We considered four mutations (denoted C, I, N and S, corresponding to mutations C59R, I164L, N51I and S108N in the experimental system, Fig. 4A), such that any genotype can be specified by a combination of mutations encoded in a vector **x** = (*x*_C_, *x*_I_, *x*_N_, *x*_S_). The term *x_i_* is 0 if mutation *i* is absent, and 1 if it is present. In this model, the fitness *f* (**x**) of any genotype **x** is given by an additive contribution from each mutation, plus two epistasis terms corresponding to pairwise interactions between C-N and C-S:

**Fig. 4.**
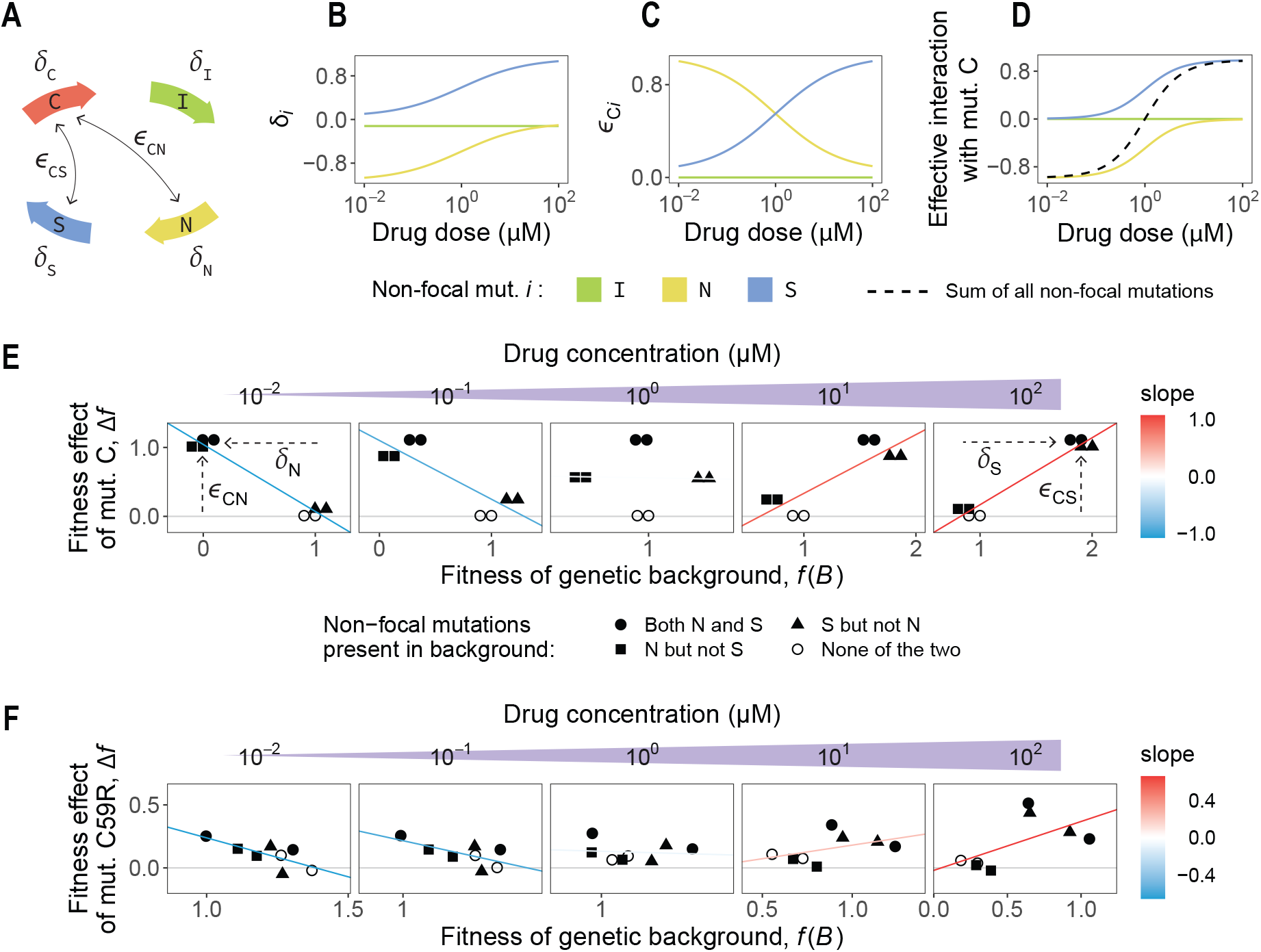
A minimal model for the environmental modulation of global epistasis. (**A**) We considered a simple model in which four mutations (C, I, N and S) have additive effects (*δ*) on fitness, and there are two pairwise interactions (*ϵ*) between C-N and C-S. (**B-C**) We made model parameters vary with environmental drug concentration as indicated. (**D**) This parameters give rise to effective interactions which mimic those observed in the empirical data. (**E**) Global epistasis slope for the focal mutation C is negative (dominated by a negative effective interaction between C and N) at low drug doses, and becomes positive (dominated by a positive effective interaction between C and S) at high concentrations. (**F**) For comparison, the empirical global epistasis patterns for C59R are included here, indicating the presence/absence of mutations N51I (N) and S108N (S) in the genetic background.

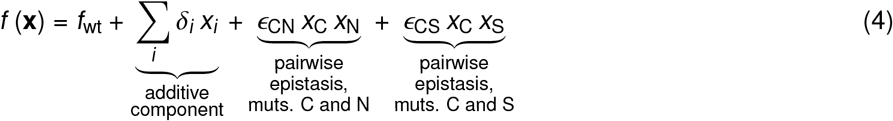

where *f*_wt_ denotes the fitness of the wild-type allele (here we take *f*_wt_ = 1 for simplicity), *δ*_*i*_ represents the (additive) fitness effect of mutation *i*, and *ϵ*_*ij*_ captures the pairwise epistasis between mutations *i* and *j*. For example, the allele carrying mutations C, I and S, **x** = (1, 1, 0, 1), has fitness equal to 1 + *δ*_C_ + *δ*_I_ + *δ*_S_ + *ϵ*_CS_. Both *δ*_*i*_ and *ϵ*_C*i*_ can in principle be positive or negative, depending on whether a specific mutation is beneficial/deleterious or a specific interaction is synergistic/antagonistic.

On top of this basic structure, we allowed the model parameters (*δ*_*i*_ and *ϵ*_C*i*_) to vary with environmental drug concentration. To reproduce the patterns observed in Fig. 3, we gave *δ*_I_ a constant value, we made *δ*_N_ increase from negative to zero with drug concentration, and we let *δ*_S_ go from zero to positive with drug dose (Fig. 4B, Materials and Methods). For simplicity, we made *δ*_C_ = 0 at all doses (note that *δ*_C_ affects the intercept, but not the slope, of the global epistasis pattern for mutation C). For the interaction parameters, we made *ϵ*_CN_ and *ϵ*_CS_ decrease and increase, respectively, with drug concentration (Fig. 4C). These choices reproduce the structure of the effective interactions observed for mutation C59R in the data (Fig. 4D, see Fig. 3C for comparison). For a drug concentration gradient between 10^−2^ μM and 10^2^ μM, we used equation 4 to quantify the fitness of each of the 16 possible combinations of mutations. We then represented the fitness effect of mutation C as a function of the genetic background fitness as previously explained in Fig. 1D. Since the model captures the structure of the effective interactions observed empirically, the global epistasis pattern for mutation C reproduces the behavior exhibited by mutation C59R in the data: the slope is negative at low drug concentrations and becomes positive at high doses (Fig. 4E).

Importantly, the minimal model allows us to interpret the change in slope sign using simple geometric reasoning. The horizontal positioning of the dots in Fig. 4E is controlled by the *δ*_*i*_ parameters, whereas the vertical positioning is dictated by the *ϵ*_C*i*_ terms. This idea is thoroughly discussed in ref. (*44*), here, for clarity, we will just analyze one example. Consider the genetic backgrounds of C, *B*(C) (where C is absent) which carry mutation N but not S (filled squares in Fig. 4E). These are, respectively, the allele carrying both I and N (**x**_1_ = (0, 1, 1, 0)), and the allele carrying N only (**x**_2_ = (0, 0, 1, 0)). Equation 4 indicates that the fitness of these genotypes, represented in the horizontal axis in Fig. 4E, is *f* (**x**_1_) = 1 + *δ*_I_ + *δ*_N_ and *f* (**x**_2_) = 1 + *δ*_N_, respectively. Since *δ*_N_ increases with drug dose (and *δ*_I_ remains invariant), these genetic backgrounds move horizontally to the right as drug concentration increases (from *f* (**x**) ∼ 0 to *f* (**x**) ∼ 1 when concentration goes from 10^−2^ μM to 10^2^ μM, Fig. 4E, leftmost and rightmost panels). Introducing mutation C in these genetic backgrounds produces the alleles **x**′_1_ = (1, 1, 1, 0) and **x**′_2_ = (1, 0, 1, 0). These have fitness *f* (**x**′_1_) = 1 + *δ*_I_ + *δ*_N_ + *ϵ*_CN_ and *f* (**x**′_2_) = 1 + *δ*_N_ + *ϵ*_CN_ respectively (remember we are setting *δ*_C_ = 0). Thus, the fitness effects of mutation C (represented in the vertical axis of Fig. 4E) are the same: Δ*f*_1_ = *f* (**x**′_1_) – *f* (**x**_1_) = *ϵ*_CN_ and Δ*f*_2_ = *f* (**x**′_2_) – *f* (**x**_2_) = *ϵ*_CN_. Because *ϵ*_CN_ decreases with drug dose, the filled squares in Fig. 4E move downwards vertically (from Δ*f* ∼ 1 to Δ*f* ∼ 0) as concentration increases.

Similar geometric reasoning can be applied to every other genetic background (hollow circles, filled circles, and filled triangles in Fig. 4E). In short, the negative slope in the leftmost panel of Fig. 4E emerges because of the horizontal negative shift produced by *δ*_N_ < 0 and the vertical positive shift produced by *ϵ*_CN_ > 0 in backgrounds carrying mutation N. The positive slope in the rightmost panel emerges because of the horizontal positive shift produced by *δ*_S_ > 0 and the vertical positive shift produced by *ϵ*_CS_ > 0 in backgrounds carrying mutation S. Thus, consistent with our reasoning, the global epistasis slope is dominated by the negative effective interaction between C and N at low drug doses, and by the positive effective interaction between C and S at high doses. This same pattern, in fact, is also observed in the empirical data when backgrounds are identified by the presence or absence of mutations N51I and S108N (Fig. 4F).

The minimal model also allows us to explore alternative situations where the global epistasis slope may be dictated by different structures in the effective interactions. For instance, in fig. S6 we show that modulation of a single interaction term can suffice for the global epistasis slope to go from negative to positive as drug concentration increases. Even in more complex scenarios, where the environment may modulate all parameters strongly, the same patterns of global epistasis can be seen as long as the effective interactions follow similar structures (fig. S7). This reinforces the idea that defining effective interactions can help rationalize the observed patterns of global epistasis and point to specific mechanisms for their environmental modulation.

## Conclusion

The observation that epistasis often has a global component has set a stepping stone in our ability to understand and predict adaptation. For instance, *diminishing returns* epistasis can explain the strikingly conserved patterns of declining adaptability reported in long-term evolution experiments (*7*–*9*,*45*,*46*). While there have been increasing efforts to understand how global epistasis patterns emerge from microscopic genetic interactions (*13*,*14*,*43*,*44*,*47*), less attention has been paid to how they might be affected by the environment. In this work, we have examined how drug concentration affects the patterns of global epistasis in a four-mutation fitness landscape. Our results show that global epistasis may be modulated by environmental variables, even ranging from *diminishing returns* to *increasing returns*. Furthermore, we have shown that the origins of this modulation can be attributed to specific gene-by-environment and gene-by-gene-by-environment interactions, making use of previous theoretical results that connected global epistasis slopes to microscopic genetic interactions (*43*,*44*). While this connection (namely equation 1) was originally formulated under the assumption of widespread epistasis (*43*), theory had suggested that it may hold even when interactions are sparse (*44*). Our results provide direct empirical evidence of equations 1 and 2 being successful at explaining global epistasis patterns even when these are dominated by a single interaction.

In particular, we have shown that the global epistasis slope of mutation C59R is dictated by interactions with two background mutations, each dominating at different drug doses. This is consistent with recent work demonstrating that global epistasis can emerge as a result of sparse genetic interactions (*14*). This, however, need not be the case for other mutations, organisms, or environmental perturbations. In fact, different focal mutations within our dataset exhibit a variety of behaviors, from a single interaction dominating throughout the entire range of drug concentrations to every interaction contributing to the emergence and shape of global epistasis. In all these cases, our work shows that defining “effective interactions”, i.e., epistatic coefficients weighted by fitness effects (equation 2) can be useful to uncover the origins for the environmental modulation of global epistasis.

Analyzing a low-dimensional landscape allowed us to directly test the fitness effect of every mutation in every genetic background, as well as every possible epistatic coefficient between all pairs of mutations. Further work will be required to determine whether effective interactions can be reliably defined and quantified in higher-dimensional landscapes, where empirically measuring the fitness of every genotype across environments might be out of reach. In addition, here we have focused on two specific drugs as our environmental variables. It will be important to test whether different forms of environmental perturbations (e.g., temperature or pH variations), and within what ranges, have the potential to modulate global epistasis. This is particularly relevant for environmental modifications induced by the population itself (*33*). More generally, our work suggests that the global epistasis framework can be readily extended to account for environmental variables, which might be critical to predict adaptive trajectories under changing environments.

## Acknowledgements

We thank members of the Sanchez and Ogbunu labs for helpful discussions.

## Funding

CBO acknowledges support from National Science Foundation’s Division of Environmental Biology Award Number 2142719.

## Author contributions

JDC, AS and CBO conceived the study. JDC analyzed and interpreted data. JDC wrote the paper, with input from AS and CBO.

## Competing Interests

The authors declare that no competing interests exist in relation to this manuscript.

## Data and materials availability

All code is available at https://github.com/jdiazc9/env_global_epist.

## Notes

https://github.com/jdiazc9/env_global_epist

### Competing Interest Statement

The authors have declared no competing interest.

