## Supplementary Material for "Environmental modulation of global epistasis is governed by effective genetic interactions"

#### Materials and Methods

**Data analysis.** Data were obtained from the original publications (*30*,*32*,*42*). All the analyses and minimal model were run using R version 4.1.2.

**Minimal model.** The basic structure of the model is specified in equation 4. Parameter values, and how they scale with environmental drug concentration (here denoted as $\lambda$ in µM units) for the model in the main text are as follows:

$$\delta_{\mathrm{C}} = 0$$
$$\delta_{\mathrm{I}} = -0.1$$
$$\delta_{\mathrm{N}} = -1 + \frac{\lambda}{1+\lambda}$$
$$\delta_{\mathrm{S}} = \frac{\lambda}{1+\lambda}$$
$$\epsilon_{\mathrm{CN}} = 1 - \frac{\lambda}{1+\lambda}$$
$$\epsilon_{\mathrm{CS}} = \frac{\lambda}{1+\lambda}$$

For the models in figs. S6 and S7, we added an interaction between mutations C and **I**, such that fitness is given by

$$f(\mathbf{x}) = f_{\mathrm{wt}} + \underbrace{\sum_i \delta_i x_i}_{\substack{\text{additive}\\\text{component}}} + \underbrace{\epsilon_{\mathrm{CI}}\, x_{\mathrm{C}}\, x_{\mathrm{I}}}_{\substack{\text{pairwise}\\\text{epistasis,}\\\text{muts. C and I}}} + \underbrace{\epsilon_{\mathrm{CN}}\, x_{\mathrm{C}}\, x_{\mathrm{N}}}_{\substack{\text{pairwise}\\\text{epistasis,}\\\text{muts. C and N}}} + \underbrace{\epsilon_{\mathrm{CS}}\, x_{\mathrm{C}}\, x_{\mathrm{S}}}_{\substack{\text{pairwise}\\\text{epistasis,}\\\text{muts. C and S}}}$$

For the first supplementary model (fig. S6), we made every parameter independent of drug concentration except for the interaction coefficient between C and S:

$$\delta_{\mathrm{C}} = 0$$
$$\delta_{\mathrm{I}} = -0.1$$
$$\delta_{\mathrm{N}} = -1$$
$$\delta_{\mathrm{S}} = 1$$
$$\epsilon_{\mathrm{CI}} = 0$$
$$\epsilon_{\mathrm{CN}} = 0.05$$
$$\epsilon_{\mathrm{CS}} = -1 + 2\,\frac{\lambda}{1+\lambda}$$

For the second supplementary model (fig. S7), we introduced more complex relationships between the parameters and the environmental variable:

$$\delta_{\mathrm{C}} = 0$$
$$\delta_{\mathrm{I}} = 2.5 - 2\,\frac{\lambda}{1+\lambda}$$
$$\delta_{\mathrm{N}} = -1.5 + 1.5\,\frac{\lambda}{1+\lambda}$$
$$\delta_{\mathrm{S}} = -2 - 0.5\,\frac{\lambda}{1+\lambda}$$
$$\epsilon_{\mathrm{CI}} = -2 + 4\,\frac{\lambda}{1+\lambda}$$
$$\epsilon_{\mathrm{CN}} = -0.5 + \frac{\lambda}{1+\lambda}$$
$$\epsilon_{\mathrm{CS}} = 1 - 2\,\frac{\lambda}{1+\lambda}$$

### Supplementary Figures

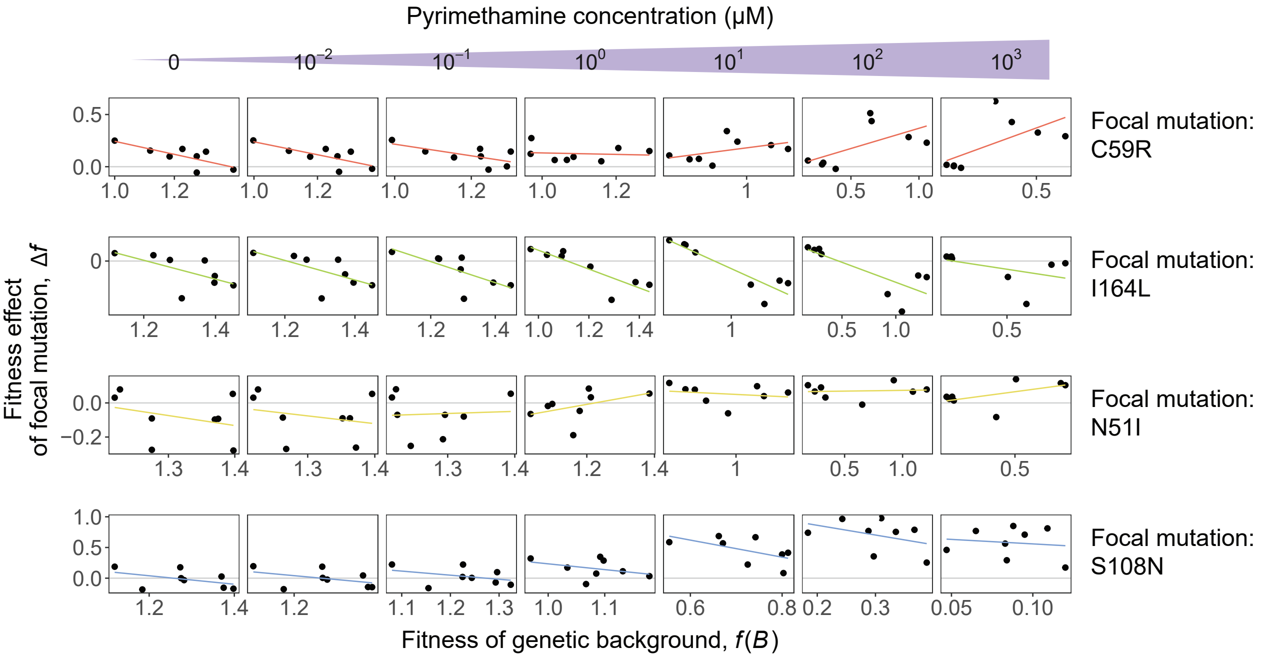


**Fig. S1. Environmental modulation of global epistasis patterns in a pyrimethamine concentration gradient.** We represent the global epistasis patterns for all four mutations (rows) in every dose of pyrimethamine from 0 to $10^3$ μM (columns). Note that the first row in this figure corresponds to Fig. 1D of the main text and is included here for completeness.

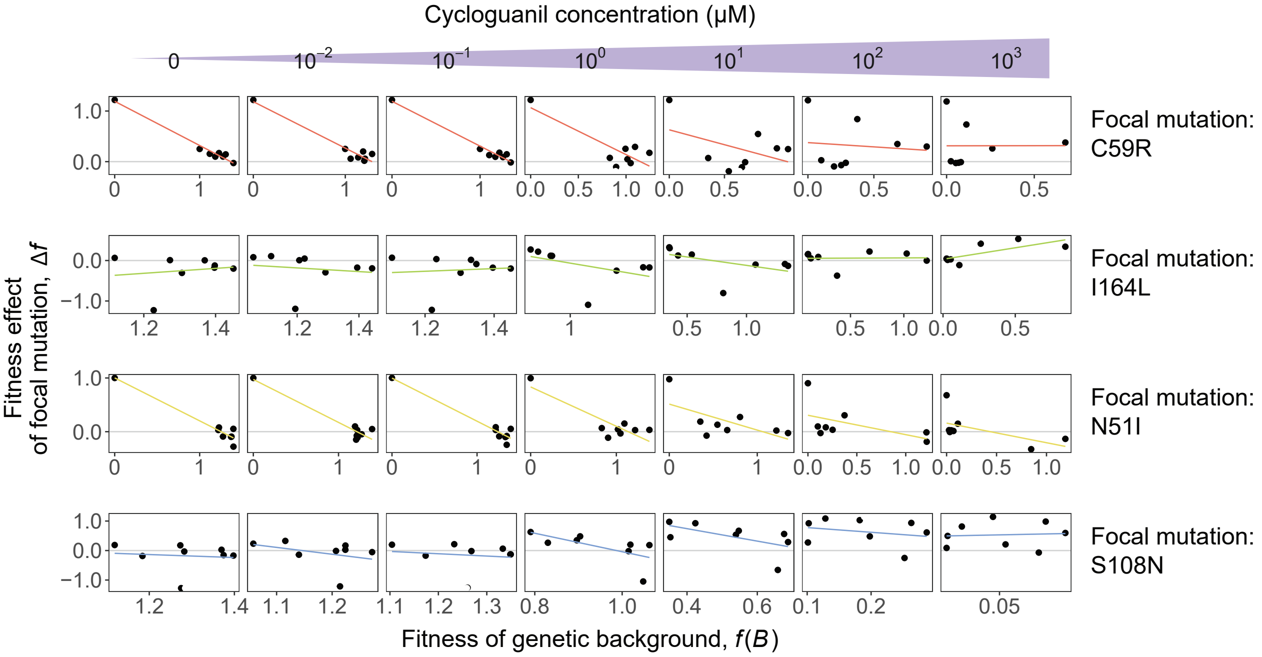


**Fig. S2. Environmental modulation of global epistasis patterns in a cycloguanil concentration gradient.** Global epistasis patterns for all four mutations (rows) in every dose of cycloguanil from 0 to $10^3$ μM (columns).

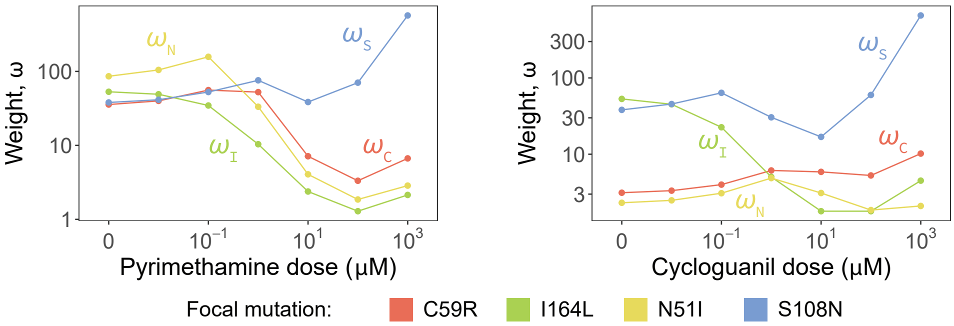


**Fig. S3. Weighting factors ($\omega$) depend on environmental drug concentration.** For each of the 4 mutations in the dataset, we represent the weighting factors $\omega_i$ ($i$ = C59R, I164L, N51I or S108N), defined in equation 3 of the main text, as a function of the drug concentration. Left panel: pyrimethamine, right panel: cycloguanil.

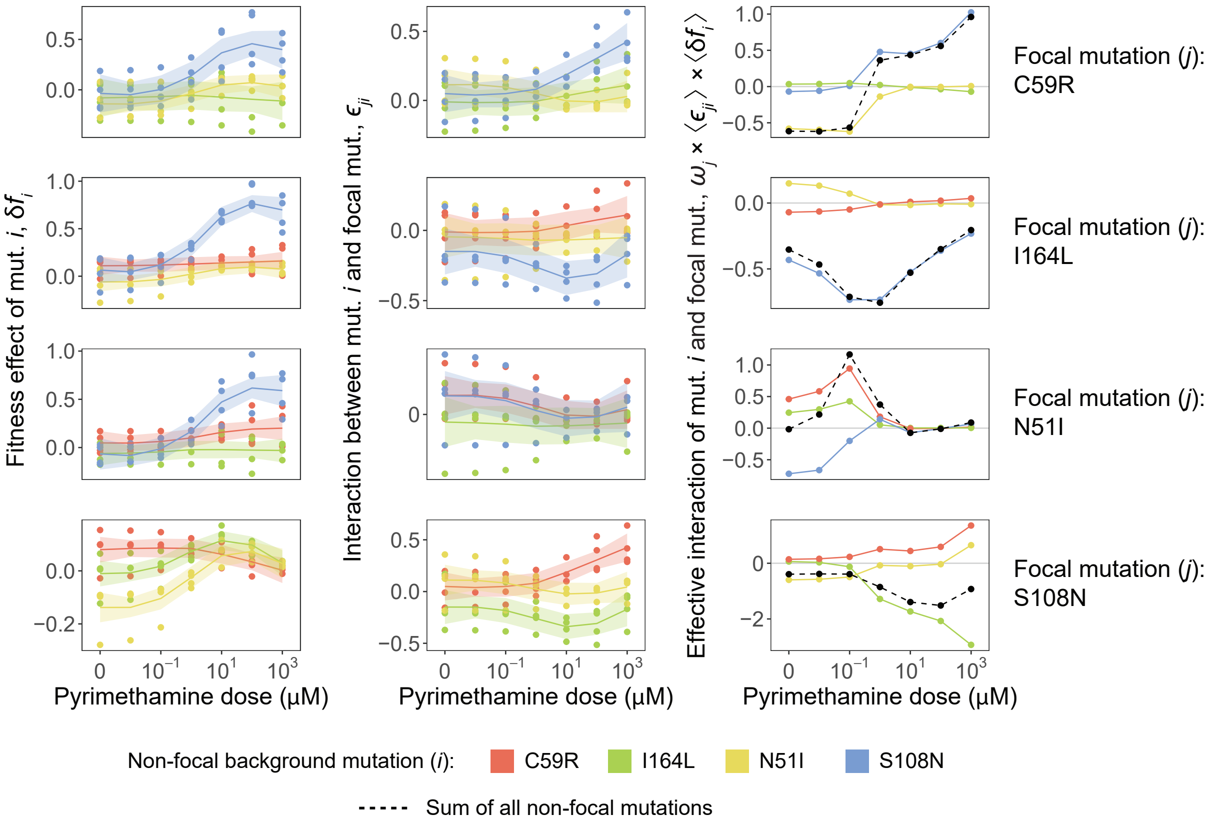


**Fig. S4. Background fitness effects and epistatic interaction coefficients for every focal mutation in a pyrimethamine concentration gradient.** Each row corresponds to a different mutation of the dataset as the focal (focal mutation is denoted as $j$). In the first column, we represent the fitness effects $\delta f_i$ of all background mutations $i$ ($i \neq j$) in every genetic background not carrying the focal mutation, $B(j)$. Each dot corresponds to a different background. In the second column, we represent the interaction terms $\epsilon_{ji}$ between the focal mutation and each of the non-focal ones. Lines are LOESS regressions and shaded areas represent 95% confidence intervals of the regressions. In the third column, we represent the effective interaction terms $\omega_\mathrm{j} \times \langle \epsilon_{ji} \rangle \times \langle \delta f_i \rangle_{B(j)}$ (as defined in equation 2 in the main text). Note that the first row of this figure corresponds to the three panels of Fig. 3 in the main text.

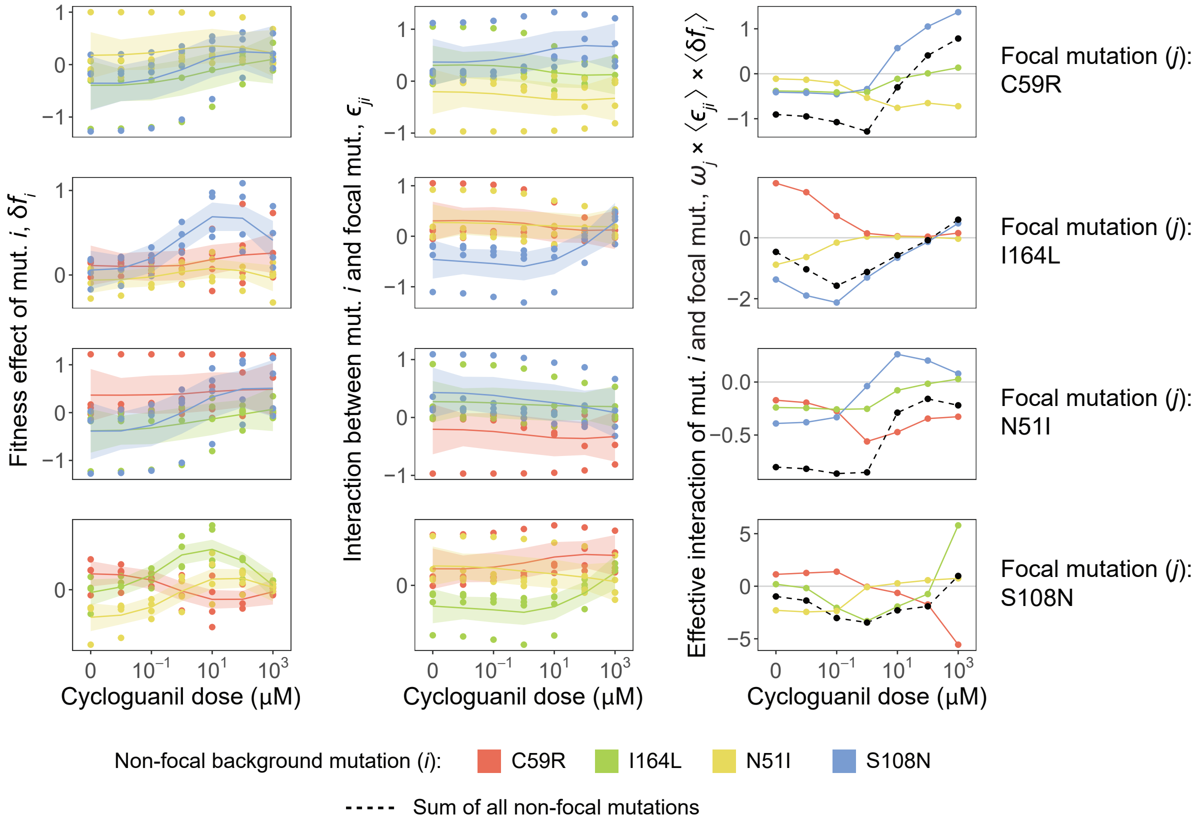


**Fig. S5. Background fitness effects and epistatic interaction coefficients for every focal mutation in a cycloguanil concentration gradient.** We represent the same information as in fig. S4, here for the cycloguanil data.

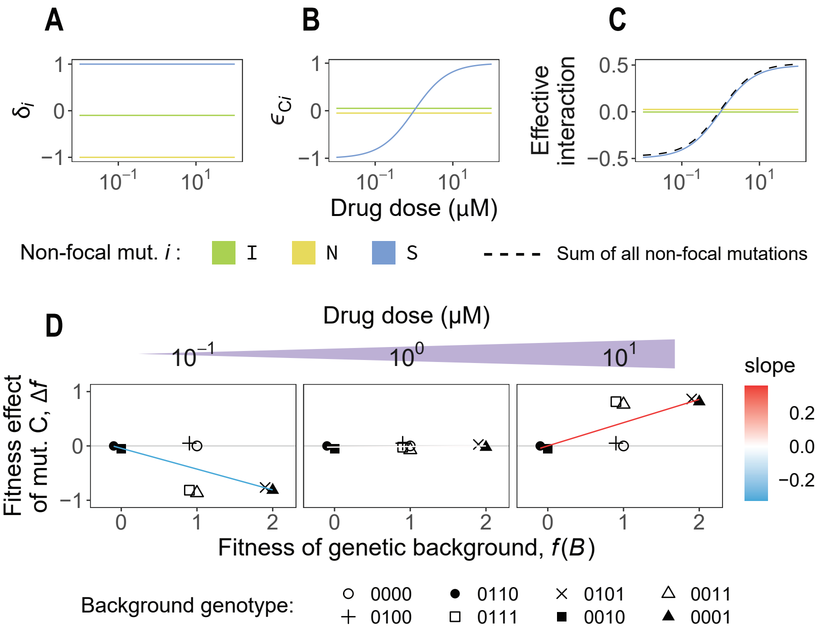


**Fig. S6. Modulation of a single interaction can induce environmental effects on global epistasis.** (**A-C**) Fitness effects, interaction terms, and effective interaction terms of the focal mutation C with every background mutation in an alternative minimal model. Here, we make it so drug concentration only modulates a single parameter, the interaction coefficient between mutations C and S ($\epsilon_{CS}$) (Materials and Methods). This interaction goes from negative to positive and dominates the sum of effective interactions (dashed line) across all drug doses. (**D**) The global epistasis slope of the focal mutation C also goes from negative to positive under these conditions.

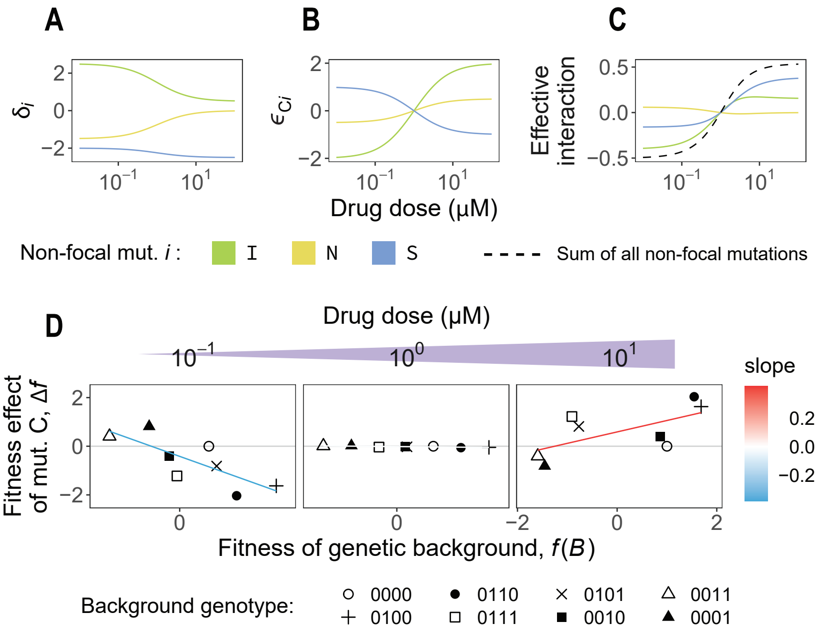


**Fig. S7. Widespread environmental effects can lead to modulation of global epistasis patterns.** (**A-C**) Fitness effects, interaction terms, and effective interaction terms of the focal mutation C with every background mutation in another alternative minimal model. In this case, we make all parameters vary strongly with environmental drug concentration (Materials and Methods). (**D**) Even here, where gene-by-environment effects are strong and widespread, effective interactions explain the environmental modulation of global epistasis.
